# An inferotemporal coding strategy robust to partial object occlusion

**DOI:** 10.1101/2024.04.09.588746

**Authors:** Andrew Cheng, Sach Sokol, Charles E. Connor

## Abstract

Object coding in primate ventral pathway cortex progresses in sparseness/compression/efficiency, from many orientation signals in V1, to fewer 2D/3D part signals in V4, to still fewer multi-part configuration signals in AIT (anterior inferotemporal cortex).^1–11^ This progression could lead to individual neurons exclusively selective for unique objects, the sparsest code for identity, especially for highly familiar, important objects.^12–18^ To test this, we trained macaque monkeys to discriminate 8 simple letter-like shapes in a match-to-sample task, a design in which one-to-one coding of letters by neurons could streamline behavior. Performance increased from chance to >80% correct over a period of weeks, after which AIT neurons showed clear learning effects, with increased selectivity for multi-part configurations within the trained alphabet shapes. But these neurons were not exclusively tuned for unique letters based on training, since their responsiveness generalized to different, non-trained shapes containing the same configurations. This multi-part configuration coding limit in AIT is not maximally sparse, but it could explain the robustness of primate vision to partial object occlusion, which is common in the natural world and problematic for computer vision. Multi-part configurations are highly diagnostic of identity, and neural signals for various partial object structures can provide different but equally sufficient evidence for whole object identity across most occlusion conditions.

## Results

We constructed letter-like shapes by crossing four top shapes with four bottom shapes (Fig. 1a). Monkeys were trained in a matching task on one of two non-overlapping 8-letter alphabets drawn from these shapes (Set 1, *green* or Set 2, *pink*). The other, held-out alphabet comprised novel letters constructed from the same, familiar elements but in different combinations. We refer to the 16 letters in the two alphabets as canonical stimuli, to distinguish them from inverted versions of the same stimuli (Fig. 1b), which served as a set of non-trained letters constructed from non-trained elements of similar complexity. Comparison of AIT responses across trained, held-out, and inverted letters allowed us to measure the effects of letter training and part training on shape coding.

**Figure 1.**
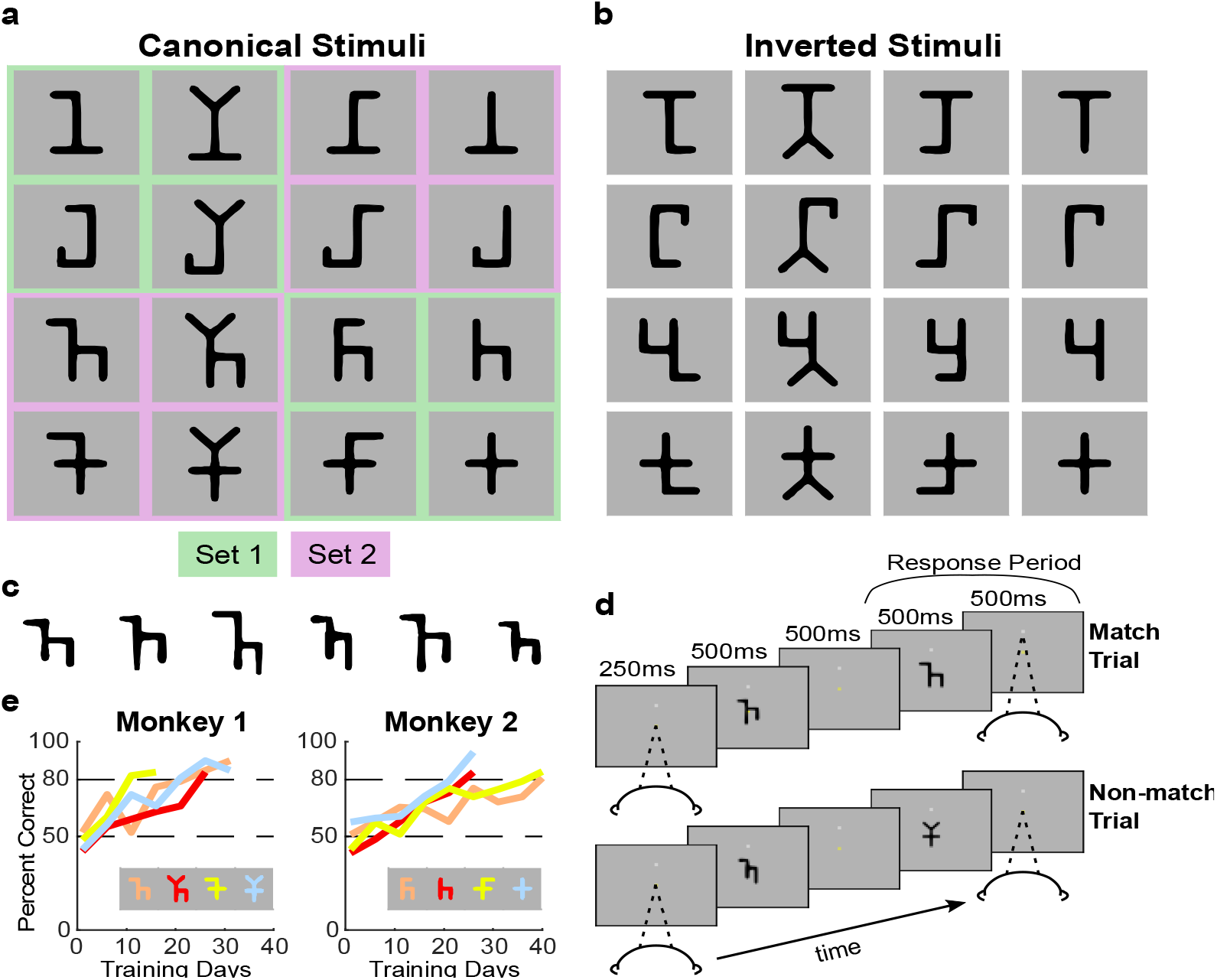
Stimuli and behavioral task. **(a)** The canonical (unmorphed) shapes in the 16 stimulus categories. Shapes were constructed by combining one of four alternative tops (*columns*) with one of four alternative bottoms (*rows*). Each monkey was trained with eight stimuli, either Set 1 (*green*) or Set 2 (*pink*). The other set served as unfamiliar shapes constructed from familiar parts. **(b)** The 16 inverted versions of the canonical stimuli, which served as unfamiliar shapes constructed from unfamiliar parts. **(c)** An example series of morphed versions of one of the canonical shapes. Individual limbs could vary in width and length. Monkeys were trained to classify these morphs as belonging to one of the canonical shape categories, requiring that they learned the general shape topology rather than any specific, local details. **(d)** The task sequence of the serial match/non-match task. The first stimulus was a morphed version of a shape category. The second stimulus was a canonical shape. The monkey earned a liquid reward by saccading upwards for a categorical match or holding fixation for a non-match. **(e)** Percent correct performance of each monkey on four categories introduced sequentially at periods of several weeks. Regardless of previous experience with earlier categories, progressing from chance (50%) to an 80% performance criterion required 20 to 30 days for most categories.

In the training task, letters were presented in a variety of morphed versions (e.g. Fig. 1c). This required monkeys to recognize the general medial axis shape of each trained letter while disregarding variations in the lengths and thicknesses of limbs. This is analogous to human recognition of letters across different fonts and handwriting styles. The task sequence (Fig. 1d) was (i) visual fixation on a center dot, (ii) sample letter, (iii) comparison letter (iv) response—hold fixation for a non-match, saccade upwards to a target for a match, (5) reward. The task was difficult due to the confusability of letters that share compositional elements and change precise shape their precise shape across different morphed versions. Training to an 80% correct criterion required weeks (Fig. 1e), consistent with previous research showing that visual learning depends on long-term changes in the selectivity and even the modular organization of neurons in high-level visual cortex.^19–29^

Following training, we recorded spiking responses of individual AIT neurons to letters during a passive fixation task. We did not measure AIT responses during the behavioral task, since our goal was only to observe shape coding as a function of familiarity, not behavioral task effects. Fig. 2A and B show responses of an example AIT neuron from an animal previously trained on alphabet 1 (*gray outlines*). This example illustrates two general findings. First, neural selectivity for individual trained letters did not increase relative to held-out letters composed of the same parts. Highest response letters were just as common in the held-out alphabet (asterisk in Fig. 2A) as in the trained alphabet. We used F ratios from one-way ANOVAs (factor=letter identity, 8 levels) to compare selectivity (response variance above noise) between the two alphabets. For this example neuron, the F ratio was greater for held-out letters (F=34.1; p=5.17e-25) than trained letters (F=2. 52; p=1.93e-2). Response sparseness, a measure of narrow selectivity, was also higher in this example for held-out (0.335) vs. trained letters (0.066).

**Figure 2.**
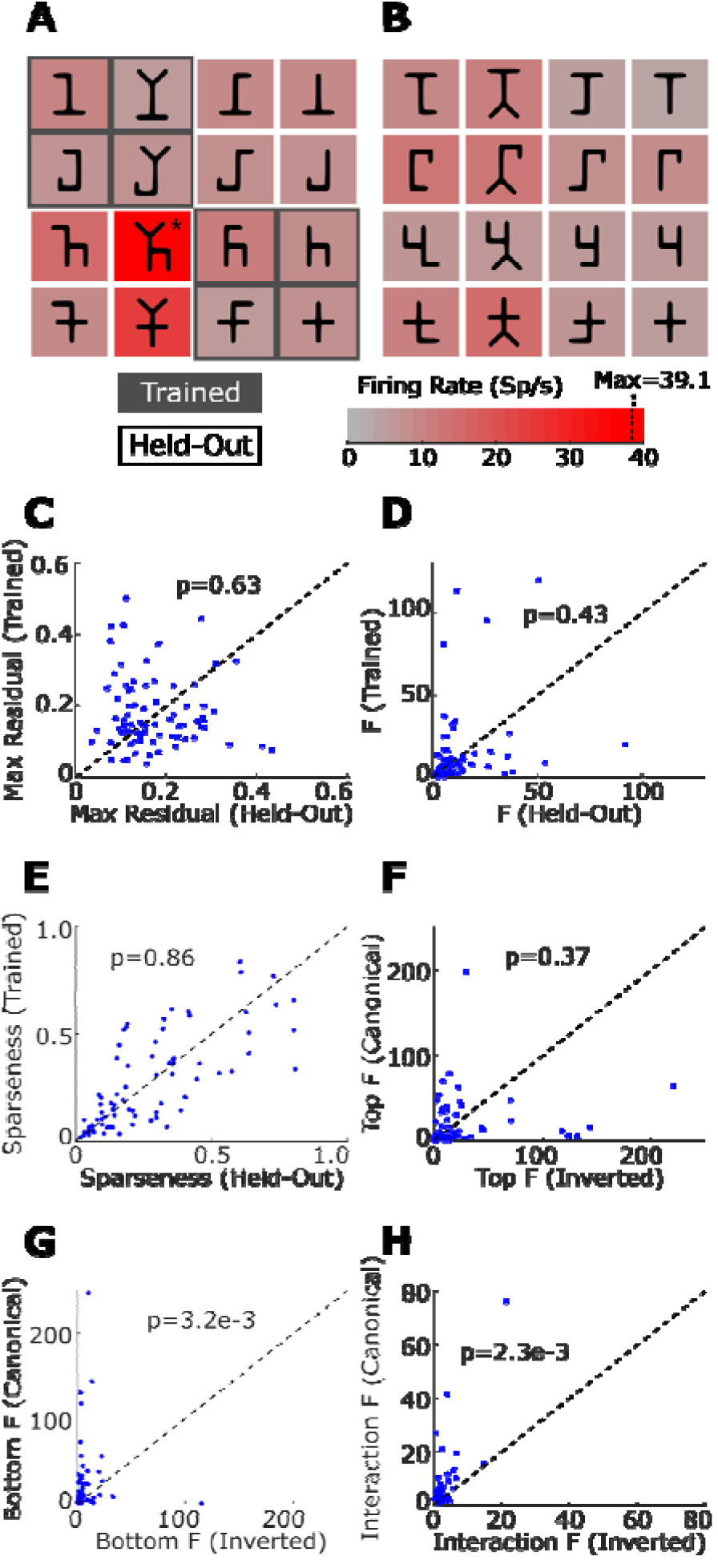
Training effects on letter responses. **(A)** Responses of an example IT neuron to canonical letters during passive fixation trials. Letters were presented in random order for 300 ms each, with 200 ms inter-stimulus intervals, for a total of 8 letters per trial and 5 repetitions per letter. This monkey was trained on alphabet 1 (*gray outlines*). The response to each letter, averaged across 300ms presentation, and across repetitions, is represented by the background color, according to the red *scale bar* at lower right. **(B)** Responses of the same neuron to the inverted letters. **(C)** Scatterplot of maximum residuals (normalized) from 2-way ANOVA of canonical letters, after subtracting main effects, comparing the trained alphabet (*vertical axis*) with the held-out alphabet (*horizontal axis*). **(D)** Scatterplot of F ratios from one-way ANOVAs performed on trained (*vertical axis*) vs. held-out (*horizontal axis*) alphabets. **(E)** Scatterplot of response sparseness for trained (*vertical axis*) vs. held-out (*horizontal axis*) alphabets. Sparseness was calculated as the inverse of response density (RD) (*23*) across eight letters:

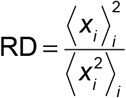

where *x*_*i*_ is the response to the *ith* object. **(F)** Scatterplot of F values top main effect in two-way ANOVA for canonical (*vertical axis*) vs. inverted (*horizontal axis*) letters. **(G)** Scatterplot of F values for bottom main effect. **(H)** Scatterplot of F values for top/bottom interaction effects.

Second, neural selectivity increased for trained partial structure in the canonical letters (partial structure was equivalent across the trained and held-out alphabets) relative to non-trained partial structure in the inverted letters. We used F ratios from two-way ANOVAs to compare selectivity for familiar (canonical) vs. unfamiliar (inverted) elements. The factors were top and bottom shape (four levels each). For the Fig. 2 example neuron, for canonical letters, main effect F ratios were 40.1 (tops; p=8.59e-21) and 37.9 (bottoms; 7.89e-20). The interaction F value, reflecting partial structure transcending the arbitrary top/bottom boundary, was 21.0 (p = 1.67e-25). For inverted letters, F ratios were only 27% (top: F=10.7, p=1.29e-6), 53% (bottom: F=20.2, p=1.31e-11), and 12% (interaction: F=2.48, p=1.03e-2) as large.

This pattern held across the population of 77 AIT neurons tested with both alphabets (41 from a monkey trained on alphabet 1, and 36 from another monkey trained on alphabet 2). For these neurons, as in the example training did not increase selectivity for familiar (trained) letters compared to novel (held-out) letters constructed from the same elements. We quantified strong selectivity for a single letter with the maximum residual response after subtracting main effects in the two-way ANOVA (e.g., 12.8 spikes/s for the example neuron). In the neural population, maximum residuals were comparable between trained and held-out alphabets (Fig. 2C; paired t-test, two-tailed, p=0.63). Overall selectivity, measured with one-way ANOVA F ratios, was also comparable between trained and held-out (Fig. 2D, p=0.43). Narrow selectivity, measured with response sparseness, was likewise similar (Fig. 2E, p=0.86).

Likewise, across a population of 57 neurons tested with both canonical and inverted letters (20 neurons from the monkey trained on alphabet 1, 37 from the monkey trained on alphabet 2), training did increase selectivity for (trained) partial structure in canonical letters compared to (non-trained) partial structure in inverted stimuli, based on two-way ANOVA. Main effects were stronger in canonical letters, though significant only for bottoms (Fig. 2F, top main effects, paired t-test, two-tailed, p=0.37; Fig. 2G, bottom main effects, p=3.24e-3). Interaction effects (transcending the top/bottom boundaries) were also greater for canonical letters (Fig. 2H, p=2.30e-3).

To explore the nature of training effects in detail, we used our previously established methods^7–11^ to analyze the geometric tuning properties of the same neural populations, based on evolving stimuli controlled by a genetic algorithm. Each test began with a first generation of 80 random letter-like stimuli, split into two independent lineages, each stimulus constructed from 1–6 limbs with random lengths, widths, orientations, and connectivity (Fig. 3A; Methods). In successive generations, higher response stimuli were more likely to produce partially morphed descendants, so that sampling in both lineages converged toward the neuron’s shape tuning range.

**Figure 3.**
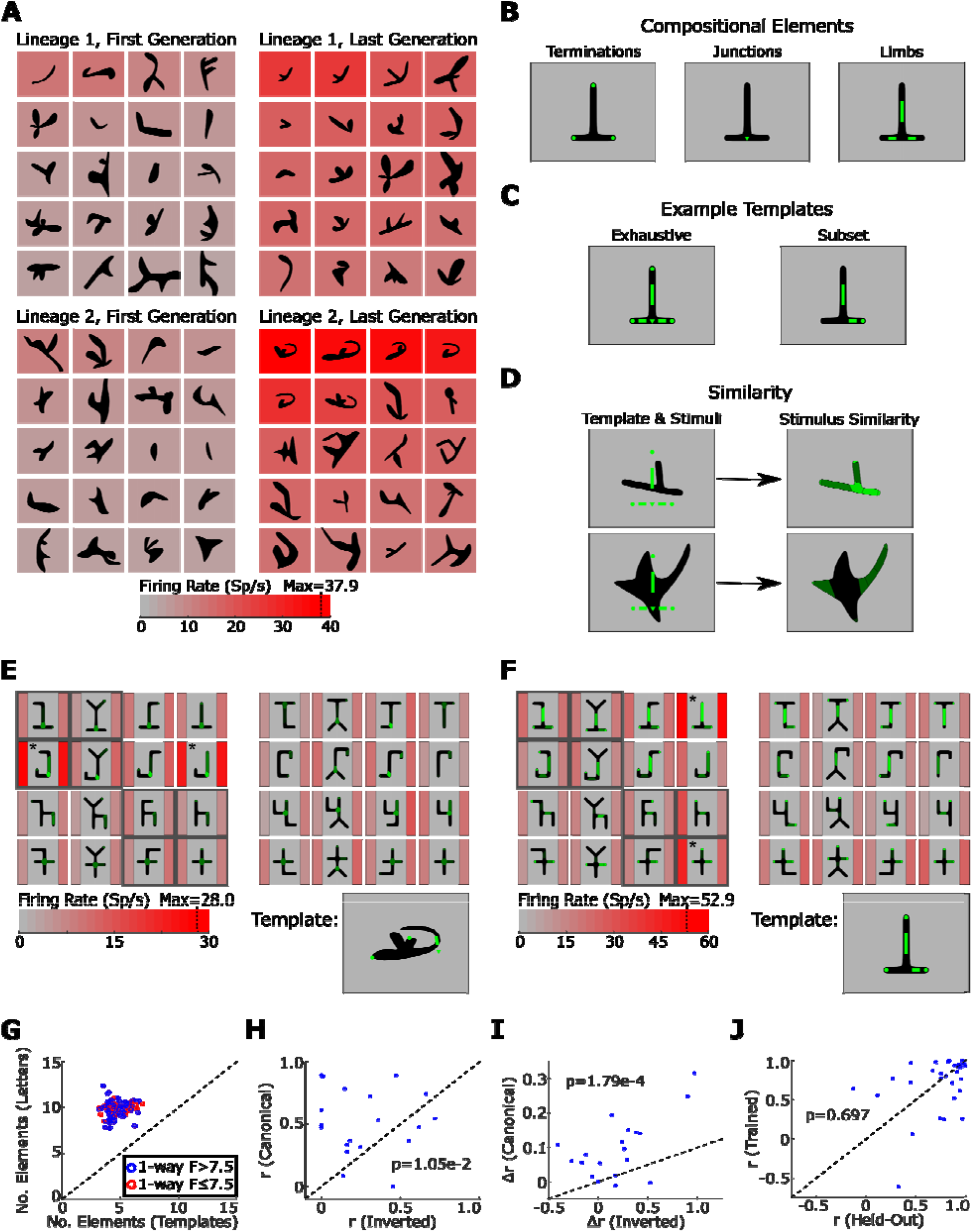
Geometric analysis of transferred shape tuning properties. (**A**) Highest response stimuli (see red *scale bar*) in first and last generations of two independent lineages of a genetic algorithm shape tuning experiment. (**B**) Examples of medial axis elements. (**C**) Examples of exhaustive and subset templates. (**D**) Element similarity between template and example shapes. (**E**) Shape tuning analysis for analysis for another example neuron. The trained alphabet is indicated by gray b(**G**) Scatterplot of number of elements in the optimum template (*x axis*) vs. a response weighted average of number of elements in the letter stimuli (*y axis*). (**H**) Scatterplot of observed-predicted correlation for inverted (*x axis*) vs. canonical (*y axis*) letters, for neurons with F_one way, trained_ > 7.5. Logistic function constants were re-fit to optimize prediction of letter responses. (**I**) Scatterplot of decrements in observed-predicted correlations due to removal of the logistic function from the template model for inverted (*x axis*) vs. canonical (*y axis*) letters, for neurons with F_one way, trained_ > 7.5. (**J**) Scatterplot of predicted-observed correlations for the held-out alphabet (*x axis*) vs. the trained alphabet (*y axis*), for neurons with F_one way, trained_ > 7.5.

Each stimulus shape was reduced to numbers describing its limbs, junctions, and terminations (Fig. 3B) in terms of position, orientation, length, and width (Methods). For each neuron, we mined high response shapes (from alphabets and genetic algorithm stimuli) for candidate tuning templates, comprising exhaustive or partial sets of elements (limbs/junctions/terminations) drawn from those shapes (Fig. 3C). We predicted responses to letter and genetic algorithm stimuli based on geometric similarity between the template and closest-match elements in the target stimulus. This is exemplified in Fig. 3D, where element similarity is indicated by *green fill* brightness for a strong match (*top*) stimulus and a weak match. The fitted parameters in the prediction model were standard deviations of Gaussian similarity functions and constants with a final logistic function to capture nonlinear tuning (Fig. S1; Methods).

Template analysis of the Fig. 3A example neuron revealed element selectivity common to the high-response trained and held-out letters (Fig. 3E). The best-fit template was based on a genetic algorithm stimulus from lineage 2. It comprised two terminations near the top and left, one vertical limb at the extreme right, and one two-limb junction at the extreme bottom right. Variations on this configuration can be seen in high response stimuli in both lineages (Fig. 3A). For each letter stimulus (Fig. 3E), similarities of closest-match elements are indicated by *green fill* brightness. Each letter is displayed with two red bars indicating observed response (*left*) and predicted response (*right*). Selectivity for the far right vertical limb and junction produced high responses (J-like letters; *asterisks*) in both trained and held-out alphabets. The vertical limb was a poor template match when it was not at the extreme right (adjacent letters in the second row). Template model predicted values correlated strongly with observed values for both trained (0.95) and held-out (0.96) letters, but not for inverted letters (0.32).

Another neuron in Fig. 3F illustrates similar element interactions in the trained and held-out alphabets. The best-fit template, derived from a held-out letter, comprised a central vertical limb, a horizontal limb at bottom right, and terminations at bottom right, bottom left, and top center. Interactions are reflected in the template analysis by the importance of the nonlinear logistic function, a soft threshold that predicts supra-linear responses more complete (multi-element) similarity. Here, the predicted/observed correlation for canonical letters was 0.81 with the logistic function and 0.49 without it, reflecting supra-linear responses to multi-element combinations in both alphabets (*asterisks*). This average correlation drop of 0.32 was similar when split out between trained (0.38) and held out (0.32) letters, but much lower for inverted letters with non-trained partial structure (0.10).

Similar results obtained across neurons. First, template tuning analyses confirmed that visual learning was confined to partial letter structure. The optimal template for each neuron comprised only a subset, approximately half (5/10), of the compositional elements in the highest response letters for that neuron (fig. 3G; paired t-test, two-tailed, p = 8.90e-84). This was equally true for neurons with stronger selectivity in the trained alphabet (F_one-way_ > 7.5; *blue circles* p = 5.87e-40) and neurons with weaker selectivity (F_one-way_ < 7.5; *red circles*) (p = 1.17e-45). Thus, multi-part shape coding is a limit on sparseness in AIT, not only for novel objects^10,11^ but also for familiar objects with high behavioral relevance.

Second, high selectivity neurons (F_one-way_ > 7.5; *blue*) exhibited stronger tuning for template elements in canonical vs. inverted stimuli. As in main effect F ratios, this was significant for elements near the bottom (Fig. 3H; paired t-test, two-tailed, p=1.05e-2) but not the top (p=0.747). Third, high selectivity neurons exhibited stronger selectivity for template element interactions in canonical vs. inverted stimuli, reflected by larger drops in observed-predicted correlations in models without logistic functions (Fig. 3I, paired t-test, two-tailed, p = 1.79e-4). Finally, enhanced selectivity for template elements and their interactions was comparable between trained alphabet letters and non-trained held-out letters, confirming that training affected multi-part selectivity but not single-shape selectivity (Fig. 3J; paired t-test, two-tailed, p = 0.697).

## Methods

Two adult male rhesus macaques (Macaca mulatta) were used for the behavioral and electrophysiology experiments. They were singly housed during training and experiments. All procedures were approved by the Johns Hopkins Animal Care and Use Committee and conformed to US National Institutes of Health and US Department of Agriculture guidelines. Both monkeys were head-restrained and trained to maintain fixation within a 0.5deg window (radius) surrounding a 0.25deg square fixation spot displayed on a monitor 60cm away. Eye positions for both eyes were monitored with a dual-camera, infra-red eye tracker (ISCAN, Inc, Woburn, MA). During behavioral training, they performed a sequential match/non-match object task (Fig. 1). During the electrophysiological experiments reported here, they performed a passive, 4 s fixation task while a total of 8 randomly selected stimuli were flashed for 300 ms each separated by 200 ms intervals. Canonical and Inverted letter stimuli were shown for at least 15 presentations per stimulus. Genetic algorithm experiments were run for neurons exhibiting responses to at least one letter stimulus two-fold greater than background spiking rate.

The electrical activity of well-isolated neurons was recorded using epoxy-coated tungsten electrodes (FHC Microsystems) and processed with a TDT RX5 Amplifier (TDT, Inc, Alachua, FL). Single electrodes were lowered through a metal guide tube via a custom-built electrode drive into the ventral bank of the superior temporal sulcus (STS), between 10 and 22 mm anterior to the auditory meatus, in the right hemispheres of both monkeys. STS was identified on the basis of structural MRI, the sequence of sulci as the electrode was lowered, and the visual response characteristics of the neurons.

For every stimulus in the genetic algorithm experiments, a medial axis skeleton was first generated (Fig. S2). Limb precursors (where each precursor is a line segment defined by the endpoint locations in two dimensions) were joined together at their endpoints to form a stimulus skeleton. Starting with a single limb precursor set to a random angle and length, a randomly generated Boolean variable decides whether to add another limb or stop the skeleton generation. An added limb will be determined whether to add another limb at each step. A maximum of six limbs per skeleton and a maximum of four limbs joined at a single node were enforced. Each skeleton was fully connected with no closed loops. The endpoints of all the limbs in the skeleton were considered nodes. From the skeleton, a surface contour was then built. Each node was assigned a width, which determined spline control points for generating the contours. For termination nodes, three or four control points (randomly) were generated about the node, producing smoother or sharper convexity respectively. For junction nodes, two or three control points (randomly) were generated for each concavity between limbs connecting to the node, again producing smoother or sharper curvatures, respectively. The resulting set of all control points defined a piece-wise B-spline contour defining the outer boundary of the shape.

This experiment depended on generating partially morphed descendants of previously tested ancestor stimuli. Morphed descendants were created by randomly altering the parameters (controlling orientations, lengths, widths, and smoothness) of a random selection of medial axis elements in the ancestor stimuli. Morphed descendants could also (randomly) undergo subtraction or addition of limbs. The first stimulus generation (80 stimuli split into two independently evolving lineages) comprised only randomly defined stimuli. Evoked responses (averaged across 5 repetitions) for these stimuli were ranked into 10 bins with equal numbers of stimuli. In the second generation, 10–20% of stimuli were randomly generated. The rest were morphed descendants of ancestor stimuli from the first generation, selected randomly, with replacement, in equal numbers from the 10 bins. In subsequent generations, ancestor stimuli were pooled across all preceding generations and re-binned.

To analyze tuning for stimulus geometry, we parameterized stimuli in terms of three types of medial axis elements: terminations, junctions, and limbs (Fig. S2A). We measured tuning in terms of templates, entire sets of subsets of medial axis elements drawn from a single letter or genetic algorithm-produced stimulus. For any stimulus in the analysis, predicted responses were based on element-wise similarity to the template:

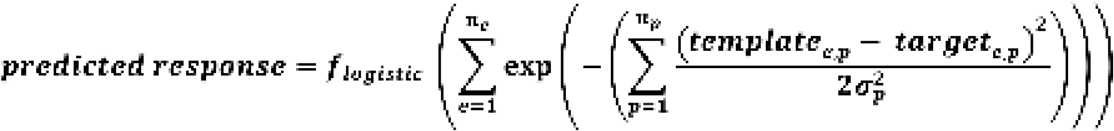

where the contribution of each parameter p (e.g. orientation) of each element e (e.g., a limb) in the template is a Gaussian function of its difference with the corresponding parameter p for the closest matching element e in the target stimulus (Fig. S2C). A combinatorial sweep of all possible one to one element matches (without replacement) of elements of the same types was tested to find the highest result for each stimulus. The logistic function parameters and standard deviations σ for each type of parameter (e.g. orientations of limbs) were fit to maximize correlation between predicted and observed responses. The logistic function captured the degree of nonlinearity of responses, i.e. the degree to which large responses depended on the combined (interactive) presence of multiple elements of the templates. Small standard deviations captured narrow, sharp tuning for a given parameter type; large standard deviations captured weak or null effects. The template producing the highest predicted/observed correlation was chosen.

Parameters for terminations were x and y position, orientation, and width. Two different types of position were considered, absolute screen position and relative position within a rectangular bounding box for the stimulus. Absolute and relative position coding were tested separately for each candidate template. Parameters for junctions were the same, but junction orientation was a vector of component limb orientations. Orientation difference between two junctions is a vector of differences between the closest match component limbs, of order equal to the number of limbs in the junction with fewer limbs. The mean and standard deviation of this difference vector were both parameters in the analysis. The mean represents an overall orientation similarity, and the standard deviation represents pairwise similarity between component limbs. Limbs were also parameterized in terms of x and y position, orientation and width, with the addition of length.

To find the template that best explained neural responses (Fig. S1), we performed a search for candidate templates from a pool of high-response stimuli, 5 to 10 stimuli from each GA lineage and 5 to 10 of the highest response letter stimuli. From this pool, an exhaustive set of all possible templates was extracted. Templates ranged from single medial axis elements to the entire shape. The resulting set of templates numbered up to roughly 40,000. From this initial set of templates, we calculated predicted firing rate without fitting standard deviations. The 1000 templates with highest correlations between predicted and observed responses were selected for further analysis. A least squares fit was used to optimize the standard deviations for each dimension for each type of parameter. The sigmoidal function was re-fitted and new predicted firing rates and correlations were assessed. Results are reported for the template models with highest correlations between predicted and observed responses.

## Acknowledgments

We thank Amy Bastian for conceptual contributions and manuscript comments. We thank Justin Killibrew, William Nash, William Quinlan, James Garmon, and Ofelia Garalde for technical assistance.

## Funding

National Institutes of Health grant R01EY029420 (AC, CEC)

National Institutes of Health grant R01NS086930 (AC, CEC)

National Institutes of Health grant R01EY025223 (AC, SS, CE)

Office of Naval Research grant N00014-20-1-2206 (AC, CEC)

## Author contributions

Conceptualization: AC, SS, CEC

Methodology: AC, SS, CEC

Investigation: AC, SS

Visualization: AC

Funding acquisition: CEC

Project administration: CEC

Supervision: CEC

Writing – original draft: AC, CEC

Writing – review & editing: AC, CEC

**Figure S1.**
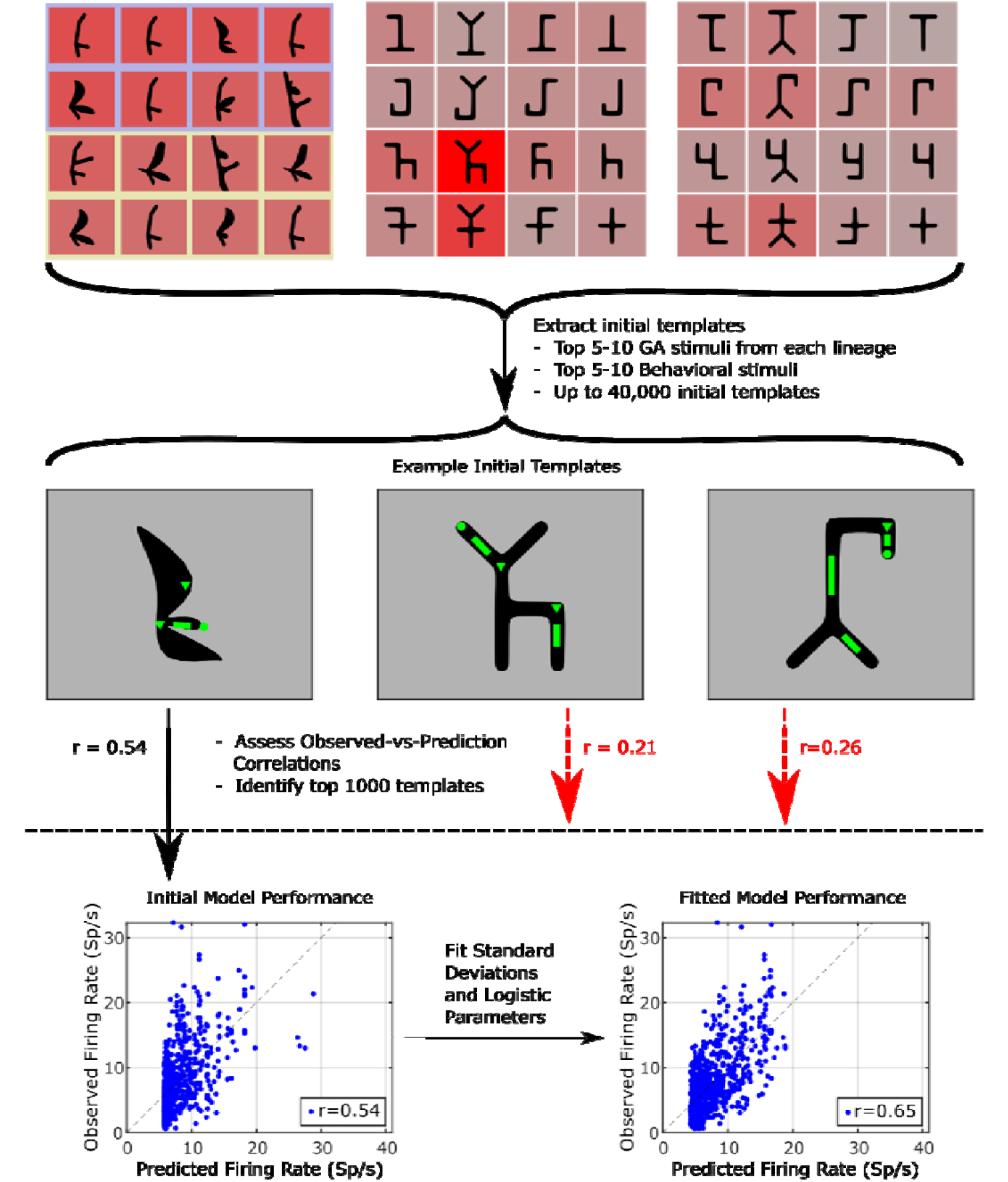
Diagram of process for selecting templates. Templates, both exhaustive and subset, were drawn from high response letter and genetic algorithm stimuli. From the resulting set of up to 40,000 templates, 1000 were selected based on highest correlation between predicted and observed responses for all stimuli prior to fitting standard deviations for medial axis parameters. For these 1000 templates, standard deviations were fit, and logistic function constants were re-fit, predicted/observed correlations were recalculated, and the highest correlation template model was selected.

## Notes

### Competing Interest Statement

The authors have declared no competing interest.

## References

1. D. J. Felleman, & D. E. Van Essen, Distributed hierarchical processing in the primate cerebral cortex. Cereb. Cortex 1, 1–47 (1991).

2. Hubel DH, Wiesel TN. 1962. Receptive fields, binocular interaction and functional architecture in the cat’s visual cortex. J. Physiol. 160:106–54

3. Hubel DH, Wiesel TN. 1968. Receptive fields and functional architecture of monkey striate cortex. J. Physiol. 195:215–43

4. Pasupathy, A., and Connor, C.E. (1999). Responses to contour features in macaque area V4. J. Neurophysiol. 82, 2490–2502.

5. Pasupathy, A., and Connor, C.E. (2001). Shape representation in area V4: position-specific tuning for boundary conformation. J. Neurophysiol. 86, 2505–2519.

6. Pasupathy, A., and Connor, C.E. (2002). Population coding of shape in area V4. Nat. Neurosci. 5, 1332–1338.

7. Srinath R, Emonds A, Wang Q, Lempel AA, Dunn-Weiss E, et al. 2021. Early emergence of solid shape coding in natural and deep network vision. Curr. Biol. 31:51–65

8. Brincat SL, Connor CE. 2004. Underlying principles of visual shape selectivity in posterior inferotemporal cortex. Nat. Neurosci. 7:880–86

9. Brincat SL, Connor CE. 2006. Dynamic shape synthesis in posterior inferotemporal cortex. Neuron 49:17–24

10. Yamane Y, Carlson E, Bowman K, Wang Z, Connor CE. 2008. A neural code for three-dimensional object shape in macaque inferotemporal cortex. Nat. Neurosci. 11:1352–60

11. Hung C-C, Carlson ET, Connor CE. 2012. Medial axis shape coding in macaque inferotemporal cortex. Neuron 74:1099–113

12. Attneave, F. (1954). Some informational aspects of visual perception. Psychol. Rev. 61, 183–193.

13. Barlow, H.B. (1959). Sensory mechanisms, the reduction of redundancy, and intelligence. NPL Symposium on the Mechanization of Thought Process. 10, 535–539. H.M. Stationery Office, London.

14. Field, D.J. (1987). Relations between the statistics of natural images and the response properties of cortical cells. J. Opt. Soc. Am. A 4, 2379–2394.

15. Treves, A., and Rolls, E.T. (1991). What determines the capacity of autoassociative memories in the brain? Network 2, 371–397.

16. Olshausen, B.A., and Field, D.J. (1996). Emergence of simple-cell receptive field properties by learning a sparse code for natural images. Nature 381, 607–609.

17. Vinje, W.E., and Gallant, J.L. (2000). Sparse coding and decorrelation in primary visual cortex during natural vision. Science 287, 1273–1276.

18. Young, M.P., and Yamane, S. (1992). Sparse population coding of faces in the inferotemporal cortex. Science 256, 1327–1331.

19. C. I. Baker, M. Behrmann, C. R. Olson, Impact of learning on representation of parts and wholes in monkey inferotemporal cortex. Nat. Neurosci. 5, 1210–1216 (2002).

20. E. Kobatake, G. Wang, K. Tanaka, Effects of shape-discrimination training on the selectivity of inferotemporal cells in adult monkeys. J. Neurophysiol. 80, 324–330 (1998).

21. M. C. Booth, E. T. Rolls, View-invariant representations of familiar objects by neurons in the inferior temporal visual cortex. Cereb. Cortex 8, 510–523 (1998).

22. D. J. Freedman, M. Riesenhuber, T. Poggio, E. K. Miller, Experience-dependent sharpening of visual shape selectivity in inferior temporal cortex. Cereb. Cortex 16, 1631–1644 (2006).

23. N. Sigala, N. K. Logothetis, Visual categorization shapes feature selectivity in the primate temporal cortex. Nature 415, 318–320 (2002).

24. D. Sasikumar, E. Emeric, V. Stuphorn, C. E. Connor, First-pass processing of value cues in the ventral visual pathway. Curr. Biol. 28, 538–548 (2018).

25. K. Srihasam, J. L. Vincent, M. S. Livingstone, Novel domain formation reveals proto-architecture in inferotemporal cortex. Nat. Neurosci. 17, 1776–1783 (2014).

26. M. J. Arcaro, P. F. Schade, J. L. Vincent, C. R. Ponce, M. S. Livingstone, Seeing faces is necessary for face-domain formation. Nat. Neurosci. 20, 1404 (2017).

27. Z. Kourtzi, L. R. Betts, P. Sarkheil, A. E. Welchman, Distributed neural plasticity for shape learning in the human visual cortex. PLoS Biol. 3, e204 (2005).

28. S. Li, S. D. Mayhew, Z. Kourtzi, Learning shapes the representation of behavioral choice in the human brain. Neuron 62, 441–452 (2009).

29. L. Cohen, S. Lehéricy, F. Chochon, C. Lemer, S. Rivaud, S. Dehaene, Language-specific tuning of visual cortex? Functional properties of the Visual Word Form Area. Brain 125, 1054–1069 (2002).

